## Supplementary information for "Prevalence and mechanisms of evolutionary contingency in human influenza H3N2 neuraminidase"

### Supplementary Table 1. X-ray data collection and refinement statistics.

| | Mos99 NA | Bil69 NA | SD93 NA + Zanamivir |
|---|---|---|---|
| **Data collection** | | | |
| Wavelength (Å) | 0.97856 | 0.97872 | 1.12723 |
| Resolution (Å) | 1.397 | 1.537 | 1.645 |
| Resolution Range [a] | 43.458-1.397 (1.401-1.397) | 46.049-1.537 (1.563-1.540) | 76.340-1.700 (1.650-1.645) |
| Space group | I 4 2 2 | C 2 | P 4 $2_1$ 2 |
| Cell dimensions | | | |
| *a*, *b*, *c* (Å) | 136.15, 136.15, 150.77 | 116.34, 137.94, 138.25 | 107.95, 107.95, 78.62 |
| *α, β, γ* (°) | 90.00, 90.00, 90.00 | 90.00, 92.16, 90.00 | 90.00, 90.00, 90.00 |
| Total reflections | 3,233,337 | 2,400,688 | 520,061 |
| Unique reflections | 138,645 | 322,494 | 54,981 |
| Multiplicity [a] | 23.0 (14.7) | 7.5 (7.5) | 9.5 (6.6) |
| Completeness (%) [a] | 99.9 (99.2) | 100.0 (100.0) | 96.8 (97.3) |
| $<I/\sigma_I>$ [a] | 20.2 (2.2) | 12.1 (2.1) | 11.5 (2.1) |
| $R_{merge}$ (%) [a] | 10.8 (126.0) | 10.4 (98.9) | 12.7 (77.4) |
| $R_{meas}$ (%) [a] | 11.3 (130.7) | 11.1 (106.4) | 13.8 (83.4) |
| $CC_{1/2}$ [a] | 0.999 (0.674) | 0.998 (0.778) | 0.998 (0.895) |
| **Refinement** | | | |
| Resolution (Å) | 43.46-1.40 | 25.00-1.54 | 25.00-1.65 |
| No. reflections | 138,634 | 304,302 | 51,828 |
| $R_{work}^c$ / $R_{free}^d$ | 0.151/0.164 | 0.166/0.180 | 0.188/0.215 |
| No. atoms | | | |
| Protein | 3,055 | 12,156 | 3,016 |
| Water | 583 | 2,004 | 317 |
| Sugars/Inhibitor | 124 | 408 | 149 |
| *B*-factors | | | |
| Protein | 14.2 | 14.9 | 18.0 |
| Sugars/Inhibitor | 25.1 | 25.5 | 33.1 |
| Water | 30.8 | 29.8 | 26.2 |
| **RMSD from ideal geometry** | | | |
| Bond lengths (Å) | 0.007 | 0.003 | 0.005 |
| Bond angles (°) | 1.12 | 0.995 | 1.210 |
| **Ramachandran statistics (%)** | | | |
| Favored | 99.7 | 95.7 | 96.1 |
| Outliers | 0.00 | 0.00 | 0.00 |
| **PDB code** | Pending | Pending | Pending |

[a] Numbers in parentheses refer to the highest resolution shell.

[b] $R_{merge} = \Sigma |I_i - <I_i>| / \Sigma I_i$ where $I_i$ = the intensity of the *i*th reflection and $<I_i>$ = mean intensity.

[c] $R_{work} = \Sigma |F_o - F_c| / \Sigma |F_o|$, where $F_o$ and $F_c$ are the observed and calculated structure factors, respectively.

[d] $R_{free}$ was calculated as for $R_{work}$, but on a test set comprising 5% of the data excluded from refinement.

**Supplementary Table 2. Primers for SD93 combinatorial mutagenesis experiment.**

| Primer name | Sequence |
|---|---|
| SD93lib-N387K-VF | 5'-CGT ACG TCT CAC TGG TCC AAA CCT AAA TCC AAA TTG CAG-3' |
| SD93lib-VR | 5'-CGT ACG TCT CAA AGC ACT TCC ATC AGT CAT TAC TAC TGT-3' |
| SD93lib-E248G-R249K-F | 5'-CGT ACG TCT CAG CTT CAG RAA RAG CTG ATA CTA AAA TAC TAT TCA TTG AGG AGG GGA AA-3' |
| SD93lib-I265T-F | 5'-TAA AAT ACT ATT CAT TGA GGA GGG GAA AAT CGT TCA TAY TAG CCC ATT GTC AGG AAG TGC-3' |
| SD93lib-336-346-5mut-F | 5'-YAT TGC CKG RAT CCT AAC AAT GAG RAA GGG RGT CAT GGA GTG AAA GGC TGG GCC-3' |
| SD93lib-336-346-5mut-R | 5'-ACY CCC TTY CTC ATT GTT AGG ATY CMG GCA ATR GCT ACT GCT GGA GCT GTC GTT-3' |
| SD93lib-E369K-R | 5'-ACT TTG AAG GTT TCA TAA CCT GAG CGT AAC TYC TCG CTG ATC GTT CTT CCC ATC CA-3' |
| SD93lib-G381E-R | 5'-CGT ACG TCT CAC CAG CCT YCA ATG ACT TTG AAG GTT TCA TAA CCT GAG CGT-3' |
| SD93lib-recover-F | 5’-CAC TCT TTC CCT ACA CGA CGC TCT TCC GAT CTA GTA ATG ACT GAT GGA AGT GCT-3’ |
| SD93lib-recover-R | 5’-GAC TGG AGT TCA GAC GTG TGC TCT TCC GAT CTA CTT GCC TAT TTA TCT GCA ATT-3’ |

**Supplementary Table 3. Primers for Bil69 combinatorial mutagenesis experiment.**

| Primer name | Sequence |
|---|---|
| Bil69lib-VR | 5'-CGT ACG TCT CAA CTC CCA TCA GTC ATT ACT ACT GT-3' |
| Bil69lib-VF | 5'-CGT ACG TCT CAG CTC AGG TTA TGA AAC TTT CAA AG-3' |
| Bil69lib-R249K-F | 5'-CGT ACG TCT CAG AGT GCT TCA GGG ARA GCC GAT ACT AGA ATA CTA TTC ATT-3' |
| Bil69lib-D286G-R | 5'-CCT ATT AGA GCC TTT CCA GTT GTC TCT GCA GAT ACA TCT GAC GYC AGG ATA TCG AGG ATA-3' |
| Bil69lib-I302V-M307V-R | 5'-CTA TAA TCT TTC AYA TTT ATG TCT ACG AYG GGC CTA TTA GAG CCT TTC CAG TTG TCT CTG-3' |
| Bil69lib-I302V-M307V-F | 5'-CCC RTC GTA GAC ATA AAT RTG AAA GAT TAT AGC ATT GAT-3' |
| Bil69lib-D329N-K334S-1-R | 5'-CTG CAA TGG CTC TTG CTA GAT CTG TCG TYG TTT CTA GGT GTG TCG CCA AC-3' |
| Bil69lib-D329N-K334S-2-R | 5'-CTG CAA TGG CTA CTG CTA GAT CTG TCG TYG TTT CTA GGT GTG TCG CCA AC-3' |
| Bil69lib-D329N-K334S-1-F | 5'-RAC GAC AGA TCT AGC AAG AGC CAT TGC AGG AAT CCT AAC AAT GAG AGA GG-3' |
| Bil69lib-D329N-K334S-2-F | 5'-RAC GAC AGA TCT AGC AGT AGC CAT TGC AGG AAT CCT AAC AAT GAG AGA GG-3' |
| Bil69lib-N356D-R | 5'-TTG CTG ATC GTT CTT CCC ATC CAC ACG TCA TTT CCA TYG TCA AAG GCC CAG CCT-3' |
| Bil69lib-L370S-R | 5'-CGT ACG TCT CAG AGC GTR AAT CCT TGC TGA TCG TTC TTC CCA TCC ACA CGT-3' |
| Bil69lib-recover-F | 5'-CAC TCT TTC CCT ACA CGA CGC TCT TCC GAT CTA CAG TAG TAA TGA CTG ATG GGA GT-3' |
| Bil69lib-recover-R | 5'-GAC TGG AGT TCA GAC GTG TGC TCT TCC GAT CTC TTT GAA AGT TTC ATA ACC TGA GC-3' |

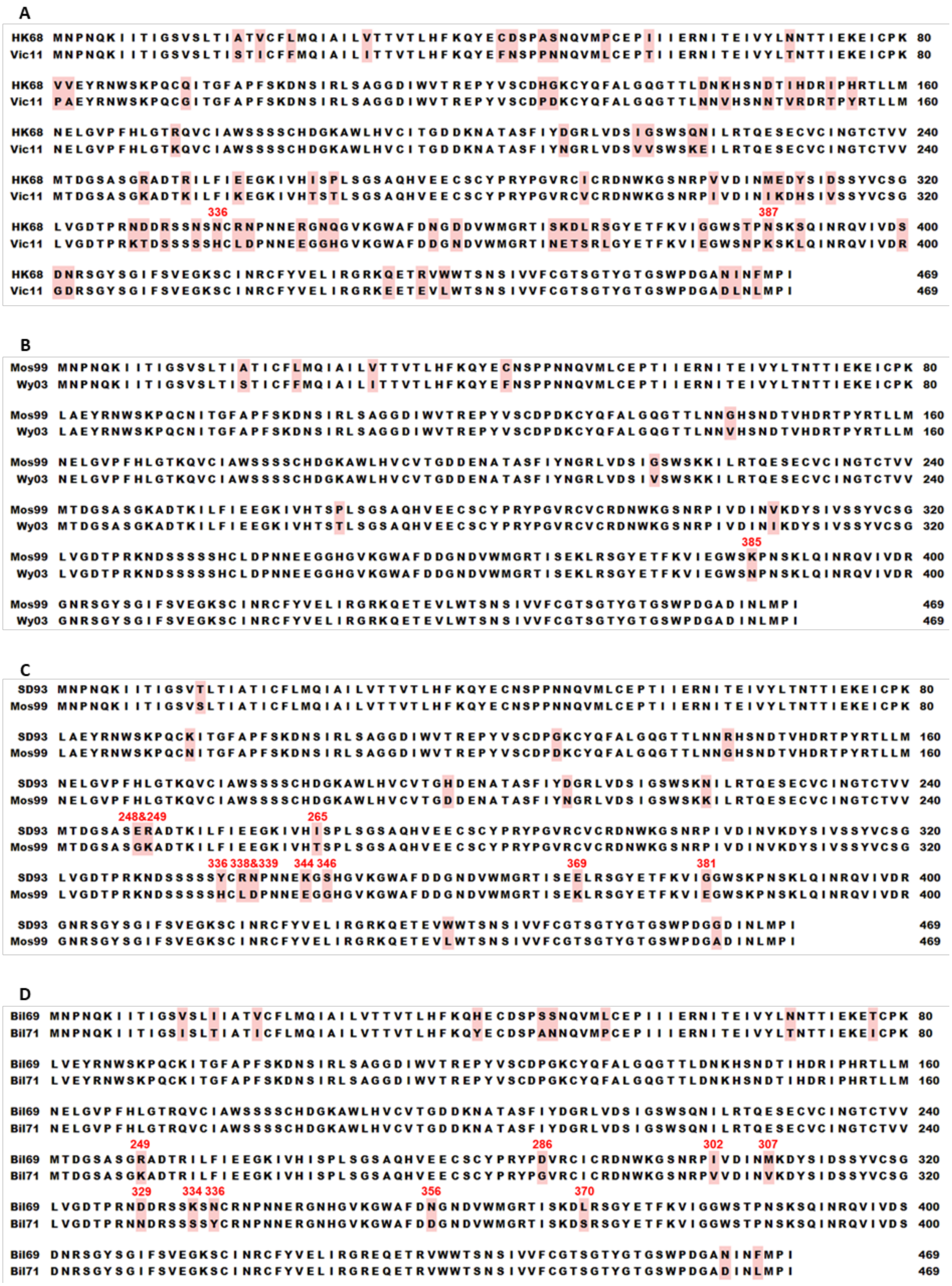


**Supplementary Figure 1. Sequence alignment of influenza virus H3N2 neuraminidase (NA).**

Alignments of NA amino acid sequences between **(A)** HK68 and Vic11, **(B)** Mos99 and Wy03 **(C)** SD93 and Mos99, and **(D)** Bil69 and Bil71. Residues that are not conserved between sequences are highlighted in pink. Mutations of interest are labeled in red. NA head domain is from residues 82 to 469.

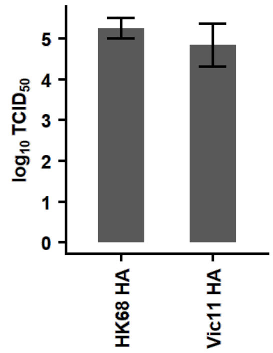


**Supplementary Figure 2. Pairing HK68 NA with Vic11 HA does not affect virus replication fitness.** The replication fitness of viruses that carry HK68 NA but different HAs (HK68 HA and Vic11 HA) was examined by a virus rescue experiment. Error bars indicate the standard deviation of three independent experiments. Virus titer was measured by $TCID_{50}$.

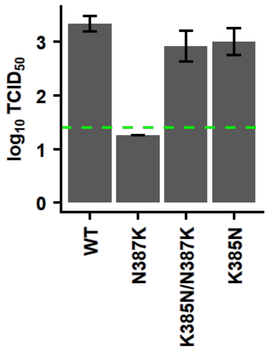


**Supplementary Figure 3. K385N of Mos99 NA does not alter virus replication fitness.** The replication fitness of different Mos99 NA mutants was examined by a virus rescue experiment. Error bars indicate the standard deviation of three independent experiments. Virus titer was measured by $TCID_{50}$. The green dashed line represents the lower detection limit.

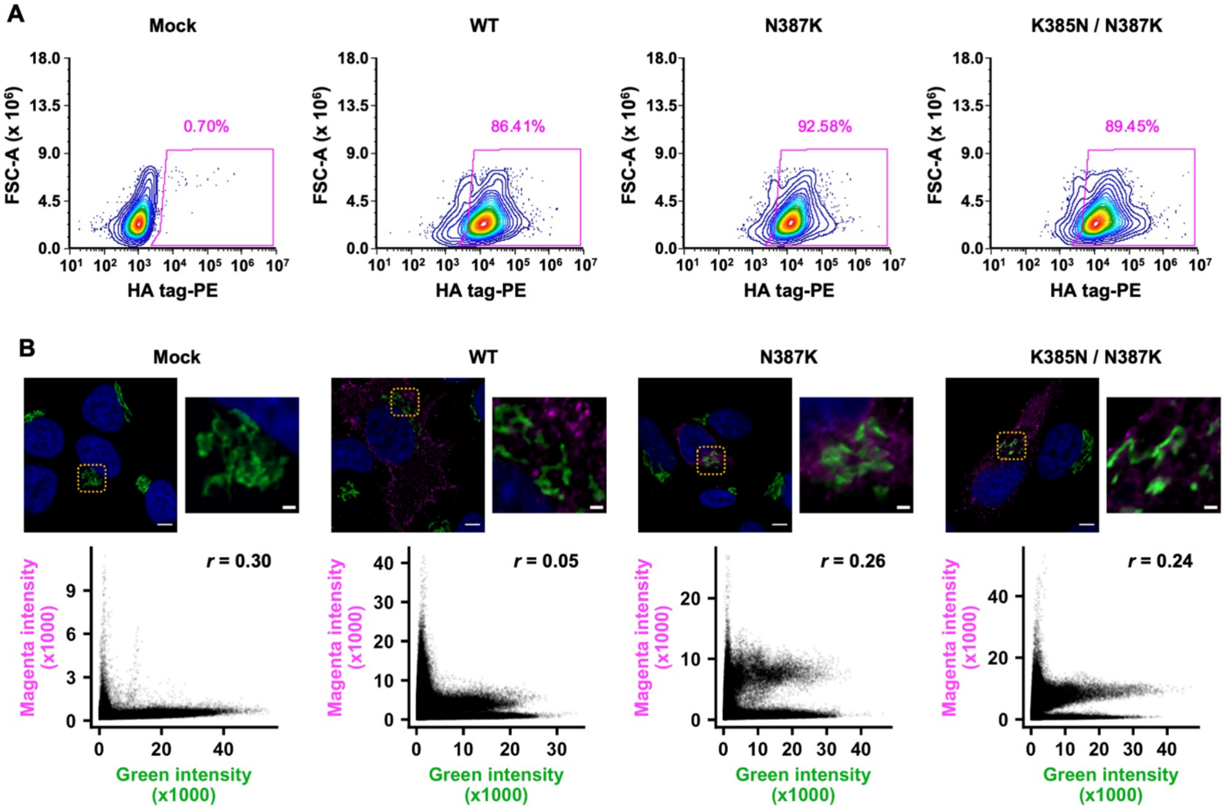


**Supplementary Figure 4. Cellular expression and localization of Mos99 NA mutants. (A)** Flow cytometry analysis of 293T cells that transiently expressed HA-tagged Mos99 NA mutants. **(B)** Confocal microscopy analysis of HA-tagged Mos99 NA mutants. Blue (DAPI), Green (GM130, Golgi), Magenta (NA). The orange box highlights the zoomed-in region, which is shown on the left. Scale bar for large image is 5 μm and scale bar for zoomed-in image is 2 μm. Below each micrograph is a cytofluorogram along with the Pearson correlation coefficient.

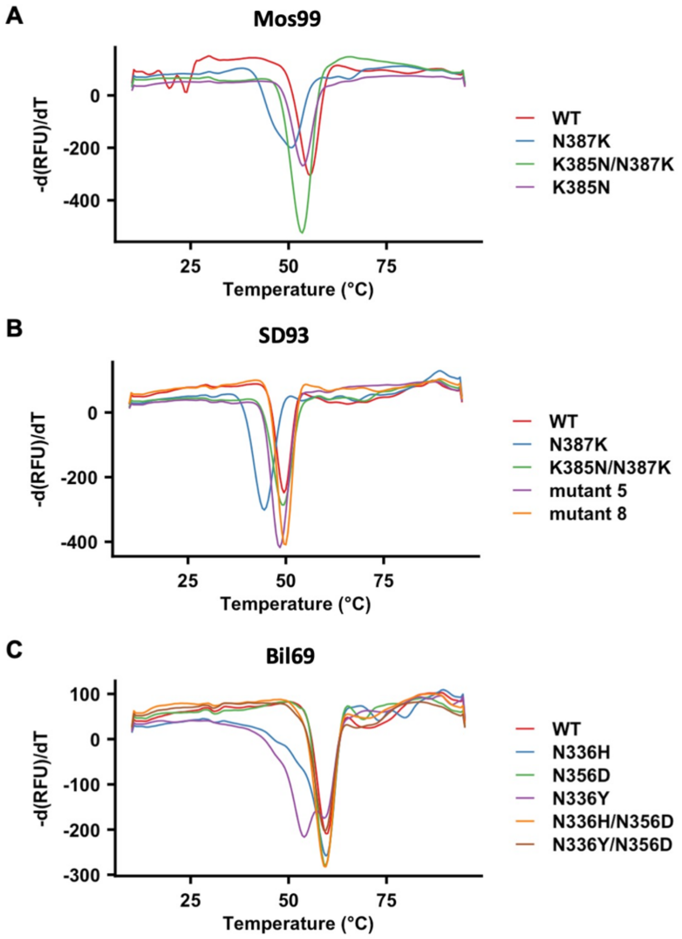


**Supplementary Figure 5. Measuring the thermal stability of different NA mutants using SYPRO orange dye-based thermal shift assay.** The first differential curves for the relative fluorescence unit (RFU) with respect to temperature were shown for **(A)** Mos99 NA WT and mutants, **(B)** SD93 NA WT and mutants, and **(C)** Bil69 NA WT and mutants. For SD93, mutant 5 represents E248G/R249K/Y336N/K344E/E369K/N387K, whereas mutant 8 represents E248G/R249K/Y336N/K344E/E369K/K385N/N387K.

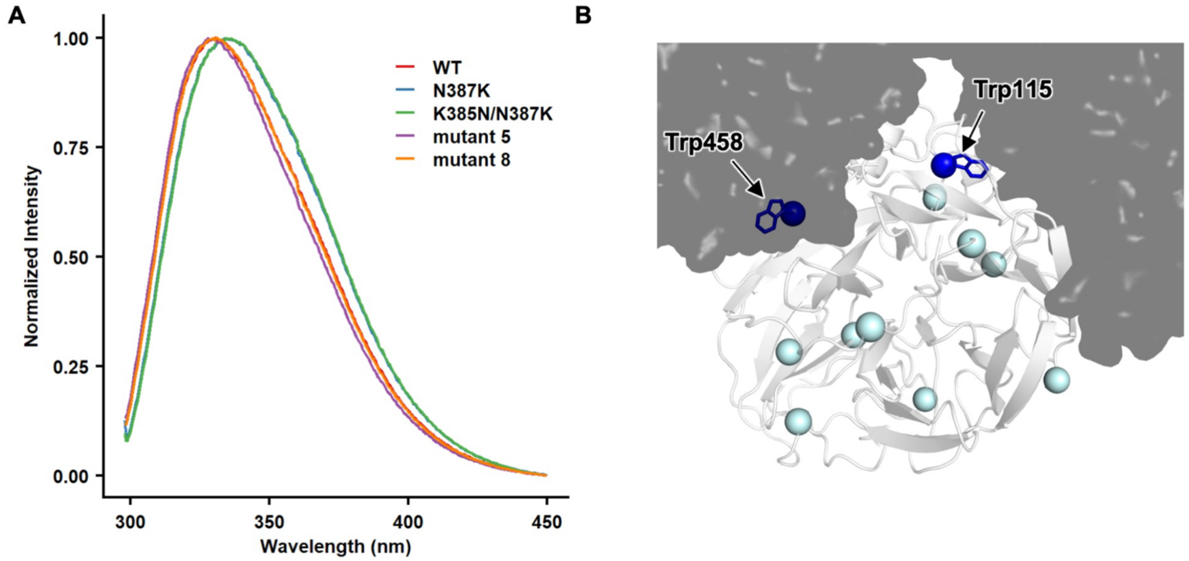


**Supplementary Figure 6. Tryptophan emission spectrum of SD93 NA WT and mutants. (A)** Normalized steady-state emission spectrum of SD93 NA WT and mutants, using $\lambda_{exc}$ = 295 nm. Of note, the blue (N387K) and green (K385N/N387K) curves almost completely overlap. Mutant 5 represents E248G/R249K/Y336N/K344E/E369K/N387K, whereas mutant 8 represents E248G/R249K/Y336N/K344E/E369K/K385N/N387K. **(B)** Two out of 12 tryptophans in the head domain of SD93 NA are located at the protomer-protomer interface. Tryptophans are shown as spheres on one protomer that is in white cartoon representation, while the other three protomers are show as semitransparent black surface. The two tryptophans at the protomer-protomer interface, namely Trp115 and Trp458, are shown in blue. Other tryptophans are in cyan.
